# SRSF2 is required for mRNA splicing and spermatogenesis

**DOI:** 10.1101/2022.09.20.508723

**Authors:** Wen-Long Lei, Zongchang Du, Tie-Gang Meng, Ruibao Su, Yuan-Yuan Li, Wenbo Liu, Si-Min Sun, Meng-Yu Liu, Yi Hou, Chun-Hui Zhang, Yaoting Gui, Heide Schatten, Zhiming Han, Chenli Liu, Zhen-Bo Wang, Wei-Ping Qian, Qing-Yuan Sun

## Abstract

RNA splicing plays significant roles in fundamental biological activities. However, our knowledge about the roles of alternative splicing and underlying mechanisms during spermatogenesis is limited. Here, we report that Serine/arginine-rich splicing factor 2 (SRSF2), also known as SC35, plays critical roles in alternative splicing and male reproduction. Male germ cell-specific deletion of *Srsf2* by *Stra8-Cre* caused complete infertility and defective spermatogenesis. Further analyses revealed that deletion of *Srsf2* disrupted differentiation and meiosis initiation of spermatogonia. Mechanistically, by combining RNA-seq data with LACE-seq data, we showed that SRSF2 regulatory networks play critical roles in several major events including reproductive development, spermatogenesis, meiotic cell cycle, synapse organization, DNA recombination, chromosome segregation, and male sex differentiation. Furthermore, SRSF2 affected expression and alternative splicing of *Stra8, Stag3* and *Atr* encoding critical factors for spermatogenesis in a direct manner. Taken together, our results demonstrate that SRSF2 has important functions in spermatogenesis and male fertility by regulating alternative splicing.

## Introduction

Spermatogenesis is a consistent and highly organized developmental process by which male germline stem cells divide and differentiate to produce mature spermatozoa. In mammalian testes, this process consists of three phases: mitosis, meiosis and spermiogenesis (1). In the first phase of spermatogenesis, mitosis is characterized by the self-renewal and differentiation of spermatogonial stem cells (SSCs), which are also known as A_single_ (A_s_) spermatogonia. There are two outlets for A_s_ spermatogonia, self-renewal to maintain the germline stem cell pool and differentiation to enter meiosis after multiple rounds of mitotic divisions of undifferentiated spermatogonia (2). A_s_ spermatogonia undergo unconventional mitotic processes to produce A_paired_ (A_pr_) spermatogonia and A_aligned_ (A_al_) spermatogonia(3). These spermatogonial progenitors including committed A_s_, A_pr_, and A_al_ spermatogonia, are uniformly identified as undifferentiated spermatogonia. Then, A_al_ spermatogonia transform into type A1 spermatogonia and further go through a series of mitoses to form A2, A3, A4, intermediate (In) and B spermatogonia. These germ cells are called differentiating spermatogonia(4). Next, B spermatogonia will divide into the pre-leptotene stage to prepare for entering meiosis which is initiated by retinoic acid (RA) and STRA8(5, 6). Any mistake in the proliferation and differentiation of SSCs can lead to failure of spermatogenesis, further resulting in severe consequences including infertility (7).

Alternative splicing (AS) is one of the most important transcriptional and post-transcriptional regulatory mechanisms to enrich the amount of mRNA and protein isoforms from a single gene, and these different protein isoforms always have different structural characteristics and functions(8-10). Generally, AS occurs more frequently in highly complex organs and organisms(11-13). There are numerous AS events during many developmental processes. Recently, it has been shown that several proteins including RAN-Binding Protein 9 (RANBP9), PTB protein 2 (Ptbp2), MORF-related gene on chromosome 15 (MRG15) and Breast carcinoma amplified sequence 2 (BCAS2) play important roles in AS events during spermatogenesis (14-17), indicating the importance of AS events during spermatogenesis, however, the functional significance of AS in the testis remains ambiguous, and the roles and regulation of AS in spermatogenesis are very limited.

The serine/arginine-rich splicing factors (SRs) have an exceedingly critical role in the alternative splicing process of precursor RNAs. The SRs can identify the splicing components of precursor RNA, then recruit and assemble spliceosomes to promote or inhibit the occurrence of alternative splicing events(18). There is a substantial amount of researches indicating that SRs are involved in nearly every step of spliceosome assembly, genomic stability, mRNA export, mRNA stability and translation(19, 20). Serine/arginine-rich splicing factor 2 (SRSF2), also known as SC35, is a member of the SRs protein family. It is an essential element of the nuclear structure, speckles(21). Recently, several studies have suggested that SRSF2 plays important roles in regulating gene transcription, mRNA stability, genomic stability, and translation(22-25). Also, some findings suggested that SRSF2 may serve as a therapeutic target for various diseases(26-29). SRSF2 is also expressed in testis, however, its functions in male germ cells are still completely unknown. Here, by crossing *Srsf2*^*Floxed/Floxed*^ (*Srsf2*^*F/F*^) mice with *Stra8-Cre* mice to generate mutant mice with specific deletion of the *Srsf2* gene in male germ cells, we found that the SRSF2 knockout caused complete infertility and germ cells were drastically lost during spermatogenesis. Further investigation revealed that deletion of the *Srsf2* gene in germ cells affected the differentiation of spermatogonia and meiosis initiation. By combining advanced linear amplification of complementary DNA ends and sequencing (LACE-seq) and RNA-seq with bioinformatics analysis, we unbiasedly mapped the binding sites of SRSF2 at single-nucleotide resolution and revealed the changes of the transcriptome and transcripts splicing in SRSF2-null testes. Our results showed that SRSF2 deletion caused abnormal alternative splicing during spermatogenesis. In particular, we found that SRSF2 directly regulated the expressions of *Stra8, Stag3* and *Atr* via AS, which have pivotal roles during spermatogenesis.

## Results

### SRSF2 is essential for male fertility

To investigate the function of SRSF2 in spermatogenesis, we first analyzed the expression of SRSF2 in the testis by using the anti-SRSF2 antibody. As a well-known marker of nuclear speckles, staining of cross-sections of seminiferous tubules in the adult mouse testis showed that SRSF2 was expressed in both germ cells and somatic cells of the testis (Figure 1A), suggesting that SRSF2 may play a potential role in spermatogenesis. Then, we generated *Srsf2* conditional knockout mice (referred to as *Srsf2*^*cKO*^) by crossing *Srsf2*^*Floxed/Floxed*^ (*Srsf2*^*F/F*^) mice in which the first and second exons were floxed (30), and *Stra8-Cre* mice in which cre activity is initiated at 3 days after birth(31). *Srsf2* was specifically deleted (Figure 1B), and the knockout efficiency of SRSF2 was confirmed by using Western blotting. The protein level of SRSF2 was significantly decreased in testes of *Srsf2*^*cKO*^ mice (Figure 1C). Thus, we successfully established male germ cell-specific knockout mice for SRSF2. The breeding assays showed that the *Srsf2*^*cKO*^ male mice were completely infertile (Figure 1D and Figure 1E). Although copulatory plugs were routinely observed, no pups were obtained when adult *Srsf2*^*cKO*^ males were mated with normal fertile females.

**Figure 1.**
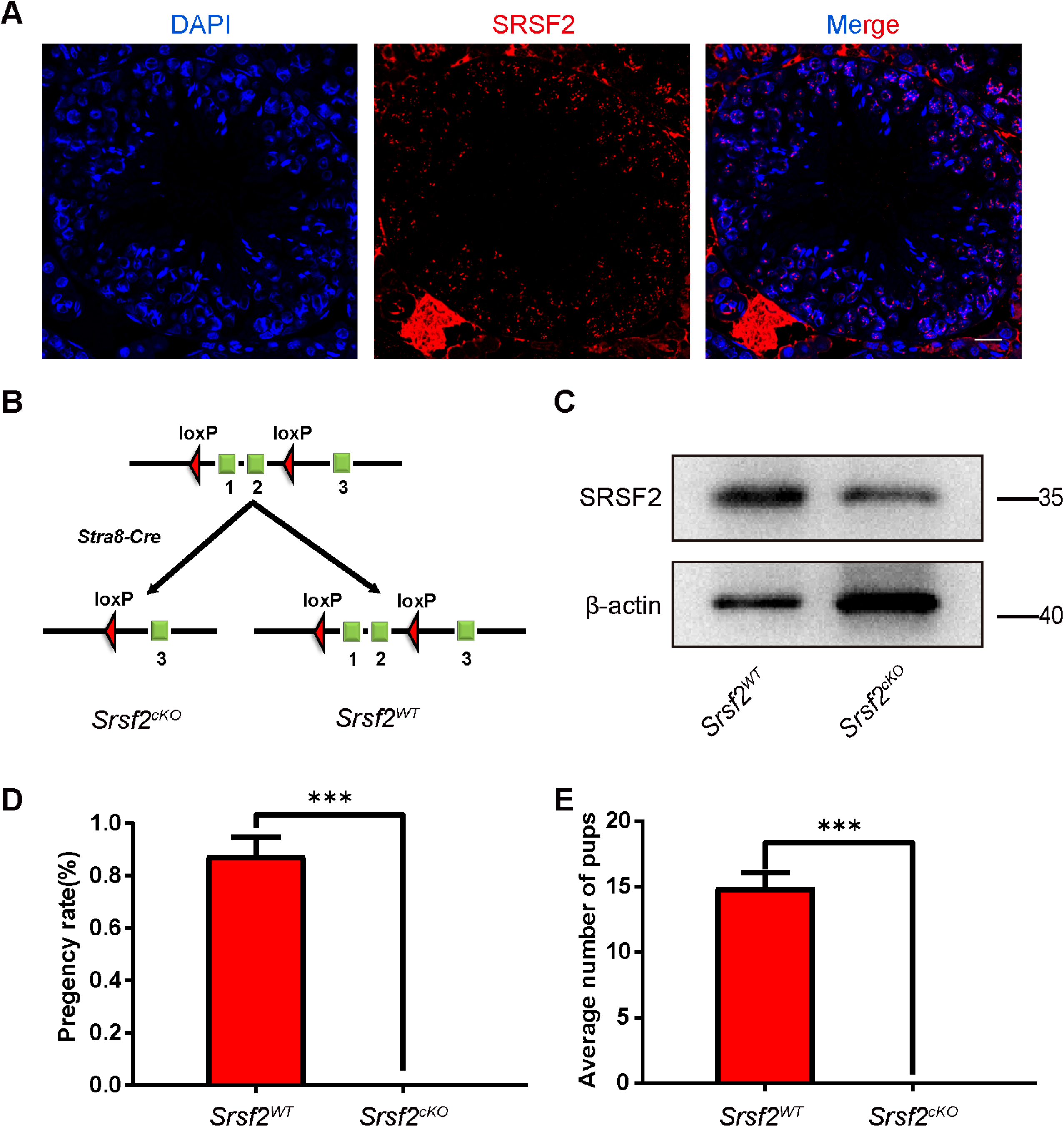
SRSF2 is essential for male fertility. (A) Representative images of localization of SRSF2 in the control testes of 8-week-old mice. The DNA was stained with DAPI (Scale bar: 20 μm). (B) Schematic diagram of deletion of *Srsf2* exons 1 and 2 and generation of *Srsf2* Δ allele by *Stra8-GFP Cre*-mediated recombination in male germ cells. (C) Western blotting analysis of SRSF2 protein in *Srsf2*^*WT*^ and *Srsf2*^*cKO*^ total testes of 8-week-old mice. β-actin was detected as an internal control. (D) Pregnancy rates (%) of plugged wild-type females after mating with *Srsf2*^*WT*^ and *Srsf2*^*cKO*^ 8-week-old males. (E) Average litter size of plugged wild-type females after mating with *Srsf2*^*WT*^ and *Srsf2*^*cKO*^ 8-week-old males. For this part, at least 3 mice (8-week-old) of each genotype were used for the analysis. Data are presented as the mean ± SEM. *P*<0.05(*), 0.01(**) or 0.001(***).

### *Srsf2* depletion causes abnormal spermatogenesis in cKO mice

To determine the reasons of infertility in *Srsf2*^*cKO*^ male mice, we firstly performed histological analyses. Compared with controls, the testes of *Srsf2*^*cKO*^ mice were much smaller (Figure 2A). The testis weight and the testis weight to body weight ratio of *Srsf2*^*cKO*^ mice was significantly lower (Figure 2B and 2C). Then we analyzed the histology of the epididymes and testes by Hematoxylin and Eosin (H&E) staining. The results showed that no mature spermatozoa were found in the epididymal lumens of *Srsf2*^*cKO*^ mice (Figure 2D). The seminiferous tubules of *Srsf2*^*WT*^ testes contained a basal population of spermatogonia, several types of spermatocytes and spermatids. However, germ cells were severely reduced in number, spermatocytes and spermatids were absent in the seminiferous tubules of *Srsf2*^*cKO*^ testes (Figure 2E). These results indicated that germ cell-specific *Srsf2* knockout results in spermatogenesis failure and thus male infertility.

**Figure 2.**
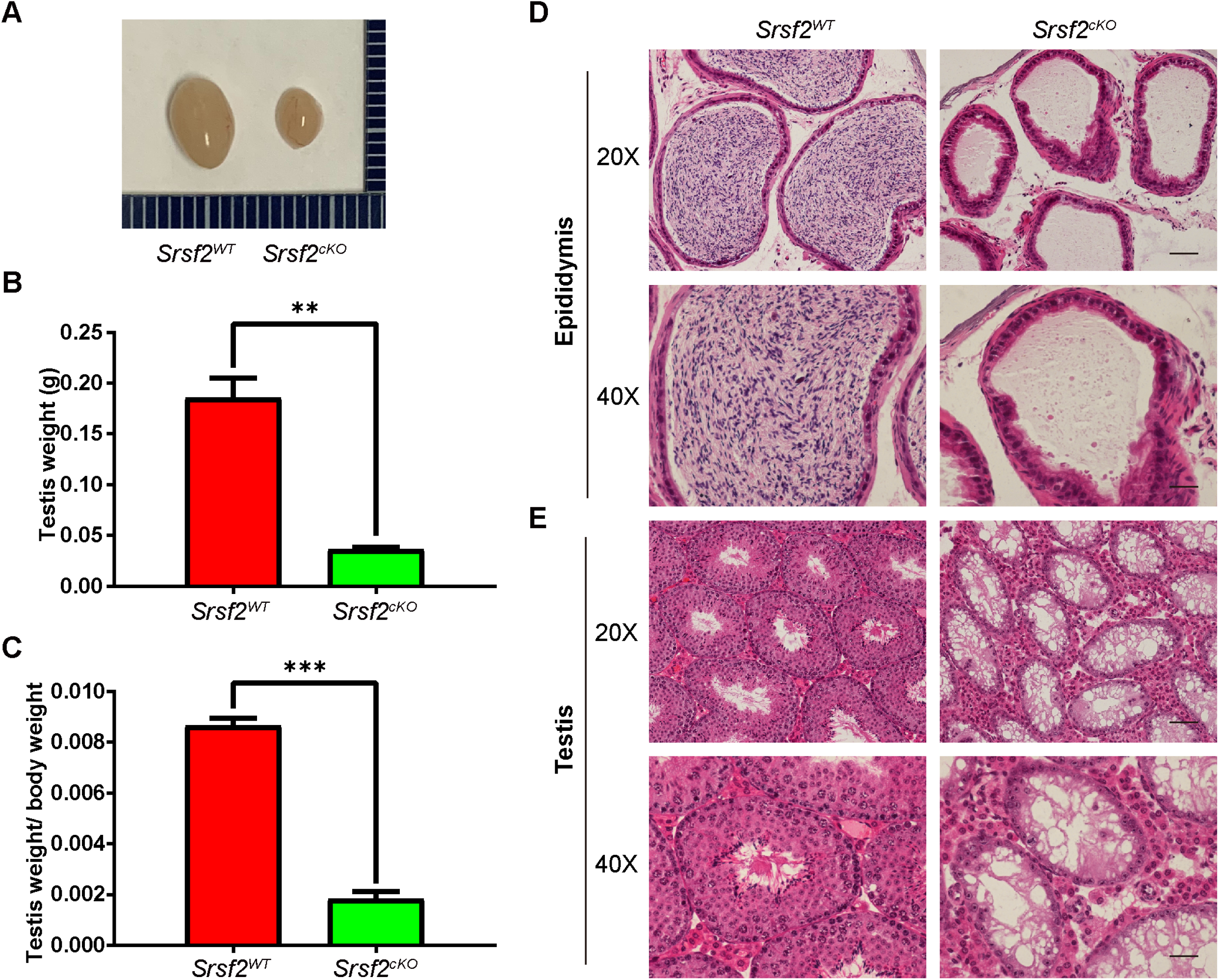
SRSF2 is required for spermatogenesis. (A) The testes of *Srsf2*^*cKO*^ were smaller than those of the control (8-week-old, the same as below). (B) Testis weight of *Srsf2*^*WT*^ and *Srsf2*^*cKO*^ 8-week-old male mice (n=3). (C) Testis weight to body weight ratio of *Srsf2*^*WT*^ and *Srsf2*^*cKO*^ 8-week-old male mice (n=3). Data are presented as the mean ± SEM. *P*<0.05(*), 0.01(**) or 0.001(***). (D) Histological analysis of the caudal epididymes of the *Srsf2*^*WT*^ and *Srsf2*^*cKO*^ mice. (Scale bar: 50 μ m) (E) Histological analysis of the seminiferous tubules of the *Srsf2*^*WT*^ and *Srsf2*^*cKO*^ mice. Scale bar: (top) 100 μm; (bottom) 50 μm.

To validate the above results, we performed immunofluorescent staining by using lectin peanut agglutinin (PNA) and antibodies against SOX9 and MVH, markers for the acrosomes of spermatids, Sertoli cells, and germ cells, respectively. Immunofluorescence results indicated that there were no PNA-positive signals in the seminiferous tubules of *Srsf2*^*cKO*^ testes and the number of MVH positive signals was significantly reduced in cKO testicular sections compared with those in the control (Figure 3A). Sertoli cells marker SOX9 staining showed that the number and location of Sertoli cells did not show an obvious change (Figure 3A).

**Figure 3.**
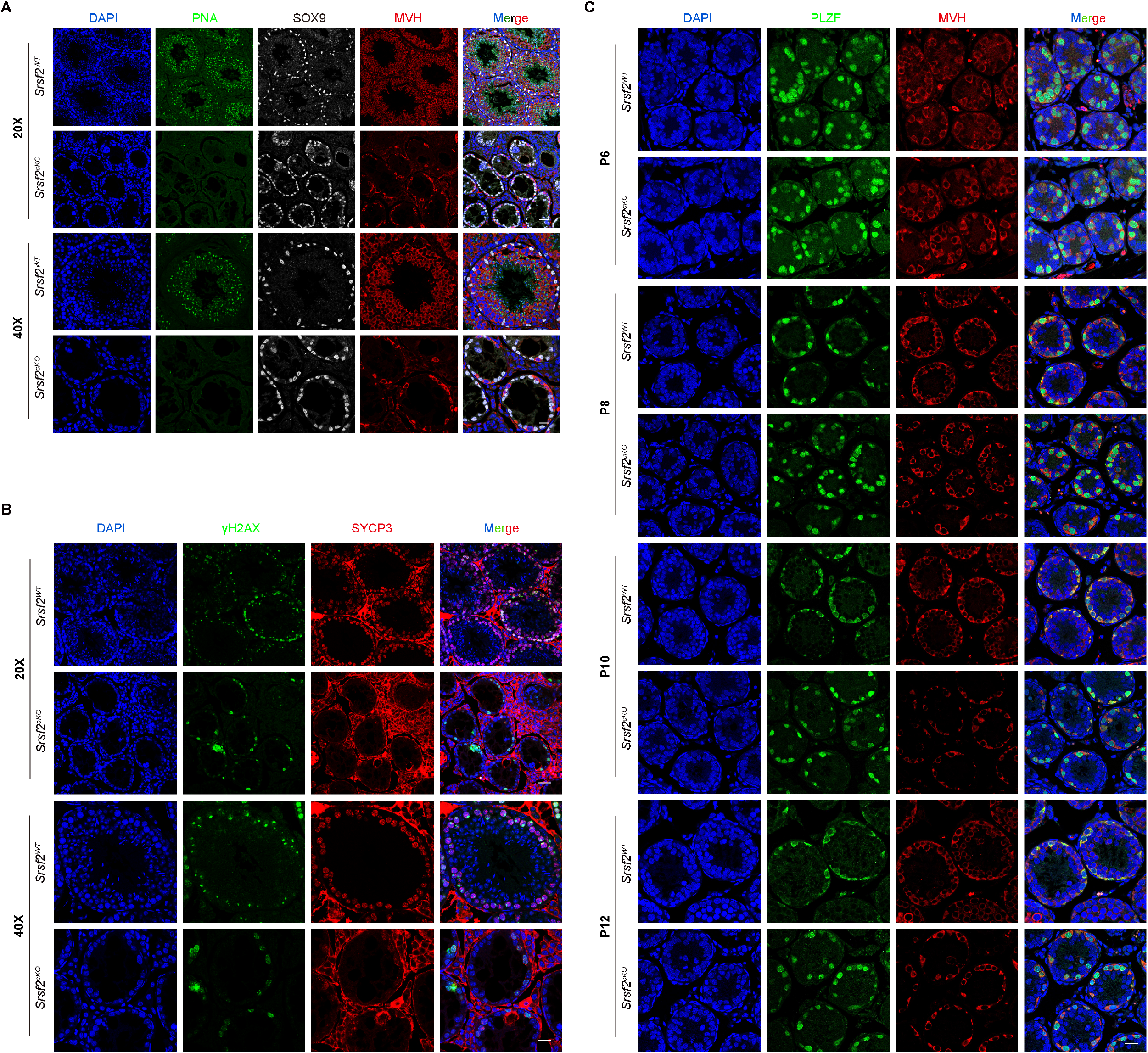
*Srsf2* deficient germ cells fail to progress into meiosis. (A) PNA-lectin histochemistry (green), SOX9 (a marker of Sertoli cells, white) and MVH (a marker of germ cells, red) immunofluorescence analysis of the *Srsf2*^*WT*^ and *Srsf2*^*cKO*^ 8-week-old male mice. Scale bar: (top) 50 μm; (bottom) 20 μm. (B) γH2AX (green) and SYCP3 (red) immunofluorescence analysis of the *Srsf2*^*WT*^ and *Srsf2*^*cKO*^ 8-week-old male mice. Scale bar: (top) 50 μm; (bottom) 20 μ m. (C) PLZF (green) and MVH (red) immunofluorescence analysis of the *Srsf2*^*WT*^ and *Srsf2*^*cKO*^ male mice at P6, P8, P10 and P12. Scale bar, 20 μm. In this part, at least 3 mice of each genotype were used for the analysis.

Meiotic recombination and homologous chromosome synapsis are two pivotal events in meiotic progression. Next we examined meiotic progression by immunostaining the axial element component of the synaptonemal complex with SYCP3 and double-strand break (DSB) marker γH2AX. Similarly, immunofluorescence results indicated that there were no SYCP3 positive signals in the seminiferous tubules of *Srsf2*^*cKO*^ testes at 8-week-old and P12, suggesting that meiosis initiation is disrupted after SRSF2 cKO (Figure 3B and Figure 3-figure supplement 1).

To further identify which stage of spermatogenesis was impaired in SRSF2-deficient mice, we performed immunofluorescence staining of the undifferentiated spermatogonia marker promyelocytic leukaemia zincfinger protein (PLZF; also known as Zbtb16) and the germ cell marker MVH (mouse vasa homologue) to characterize the first wave of spermatogenesis in mice at postnatal day 6 (P6), P8, P10, and P12. The results showed that nearly all the germ cells were undifferentiated spermatogonia in both the *Srsf2*^*WT*^ and *Srsf2*^*cKO*^ group at P6 (Figure 3C). Then the undifferentiated spermatogonia proliferated to self-renew or divided into differentiating spermatogonia from P8 to P12 in the *Srsf2*^*WT*^ group. However, MVH positive signals and PLZF positive signals were always nearly co-localized in the *Srsf2*^*cKO*^ group from P6 to P12 (Figure 3C). Altogether, these results indicated that the differentiation of spermatogonia was affected in *Srsf2*^*cKO*^ mice, which may further cause the failure of meiosis initiation.

### Changes in transcriptome and splicing of transcripts in SRSF2-null testes

According to the above-presented data, SRSF2 cKO mice displayed defects in spermatogenesis. To investigate a comprehensive perspective of the mechanisms of SRSF2 deletion in male germ cells, we isolated mRNA from *Srsf2*^*WT*^ and *Srsf2*^*cKO*^ testes at P10 and then performed RNA sequencing (RNA-seq). RNA-seq results firstly showed the reduction of *Srsf2* RNA in *Srsf2*^*cKO*^ mice testes (Figure 4A). Clustering and principal component analysis (PCA) clearly distinguished the gene expression patterns of *Srsf2*^*cKO*^ mice testes from the *Srsf2*^*WT*^ mice testes (Figure 4B). A total of 977 genes were upregulated, and 1742 genes were downregulated in *Srsf2*^*cKO*^ testes (P value of <0.05, |log2FoldChange| ≥ 0.6) (Figure 4C). Heatmap showed hierarchical clustering of differential expression genes (DEGs) of *Srsf2*^*WT*^ and *Srsf2*^*cKO*^ testes (Figure 4D). To obtain more comprehensive information, we then performed Gene Ontology (GO) annotation. GO analysis showed that these upregulated genes were involved in reproductive development, sex differentiation, and gonad development (Figure 4E). Meiotic cell cycle, chromosome segregation, DNA repair, DNA recombination, and cellular processes involved in reproduction in multicellular organisms were significantly enriched among these downregulated genes (Figure 4E). In short, these differential expression genes may account for the SRSF2-null phenotypes in spermatogenesis.

**Figure 4.**
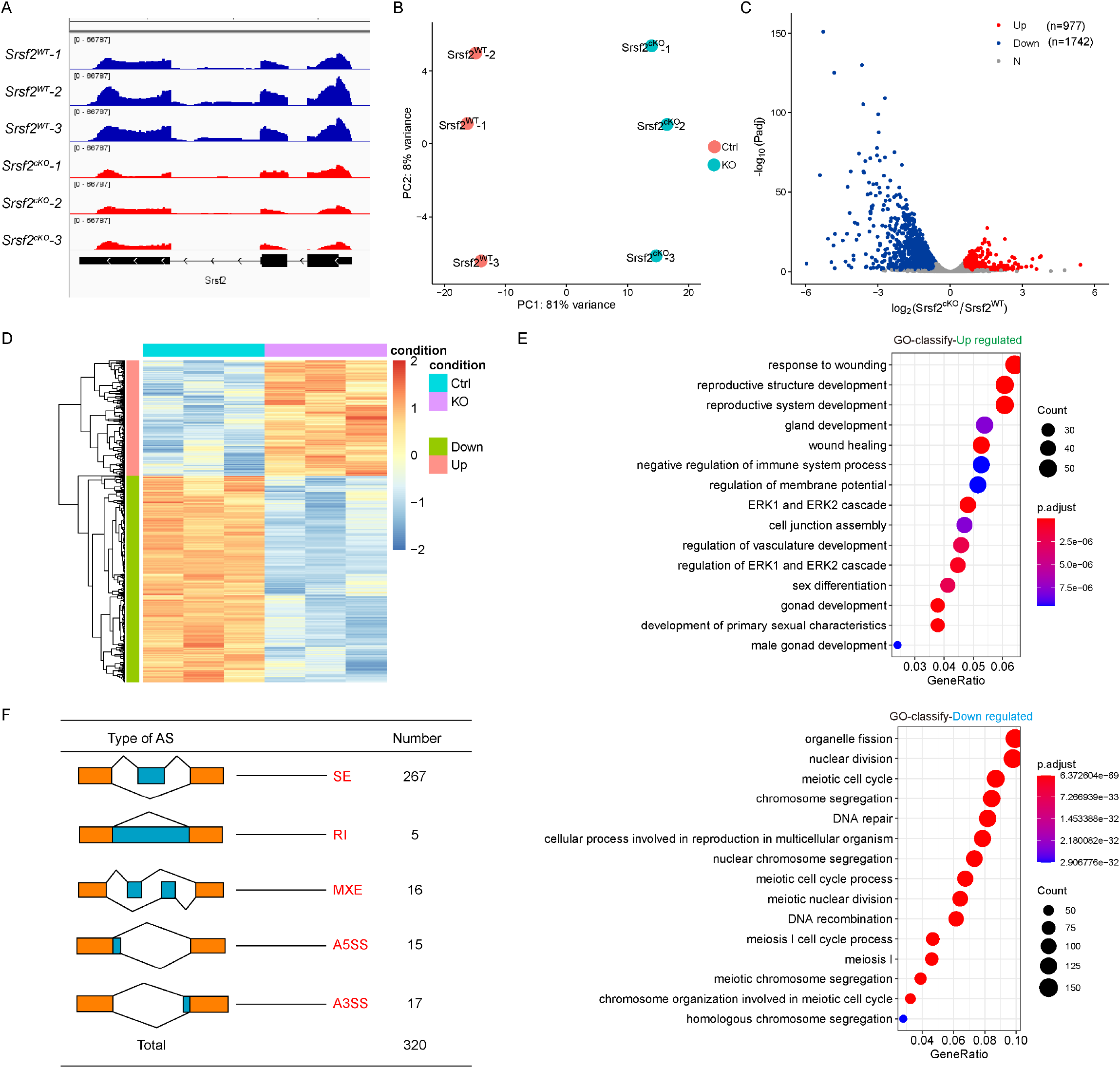
Transcriptome and splicing of transcripts changes in SRSF2-null testes. (A) RNA-seq results showing the reduction of *Srsf2* RNA in *Srsf2*^*cKO*^ mice testes. Three independent RNA-seq experiments are shown. (B) *Srsf2*^*cKO*^ groups rather than to *Srsf2*^*WT*^ groups are clustered together by PCA. (C) Volcano plot showing transcriptome changes between *Srsf2*^*WT*^ and *Srsf2*^*cKO*^ testes. (D) Heatmap showing hierarchical clustering of differential expression genes of *Srsf2*^*WT*^ and *Srsf2*^*cKO*^ male mice testes. (E) GO term enrichment analysis of upregulated genes. (F) GO term enrichment analysis of downregulated genes. (G) The five different types of alternative splicing (AS) events. The numbers of abnormal AS events were counted between *Srsf2*^*WT*^ and *Srsf2*^*cKO*^ testes by rMATS software.

Because SRSF2 played critical roles in regulating RNA splicing, we then analyzed the five different types of AS events between *Srsf2*^*WT*^ and *Srsf2*^*cKO*^ testes by using the rMATS computational tool. Compared with the *Srsf2*^*WT*^ group, a total of 320 AS events were identified as significantly changed in the *Srsf2*^*cKO*^ group (|Diff| > 0.05, FDR < 0.001). Among these 320 changed AS events, most (267) of AS events were skipped exons (SE). Moreover, there were 17 alternative 3′ splice sites (A3SS), 15 alternative 5′ splice sites (A5SS), 16 mutually exclusive exons (MXE), and 5 retained introns (RI) (Figure 4F and Figure 4-figure supplement 1). Together, these results suggested that SRSF2 is essential for RNA splicing during spermatogenesis.

### Binding landscape of SRSF2 proteins analysis in mouse testes

To further investigate the molecular mechanisms by which SRSF2 causes the failure of spermatogenesis, we performed LACE-seq analysis by using testes at P10 to profile SRSF2-binding sites in testes (Figure 5A). Two independent replicates with a high correlation in read counts were pooled together for the following analysis (Figure 5B). Among these SRSF2 clusters, more than half of them were derived from intergenic regions, while others were aligned to intron, CDS (coding sequence), UTR3 (3′ untranslated region), and UTR5 (5′ untranslated region) (Figure 5C). We also found that SRSF2 “preferentially” bound to exons and enriched between 0 and 100 nt of the 5′ and 3′ exonic sequences flanking the constitutive splice sites as revealed by analyzing the distributions of SRSF2-binding peaks within 500 nucleotides (nt) upstream or downstream of the constitutive splice site (Figure 5D). Among these SRSF2 peaks, most of them had at least one CG-rich hexamer, and more than half of the peaks contained at least one top-10 motif (Figure 5E and Supplementary file 2). GO analysis showed that these SRSF2-binding genes were involved in the regulation of RNA splicing, reproductive development, male sex differentiation, regulation of synapse organization, and regulation of chromosome segregation (Figure 5F and Figure 5G). Together, these analyses suggested that SRSF2 is essential for reproductive development.

**Figure 5.**
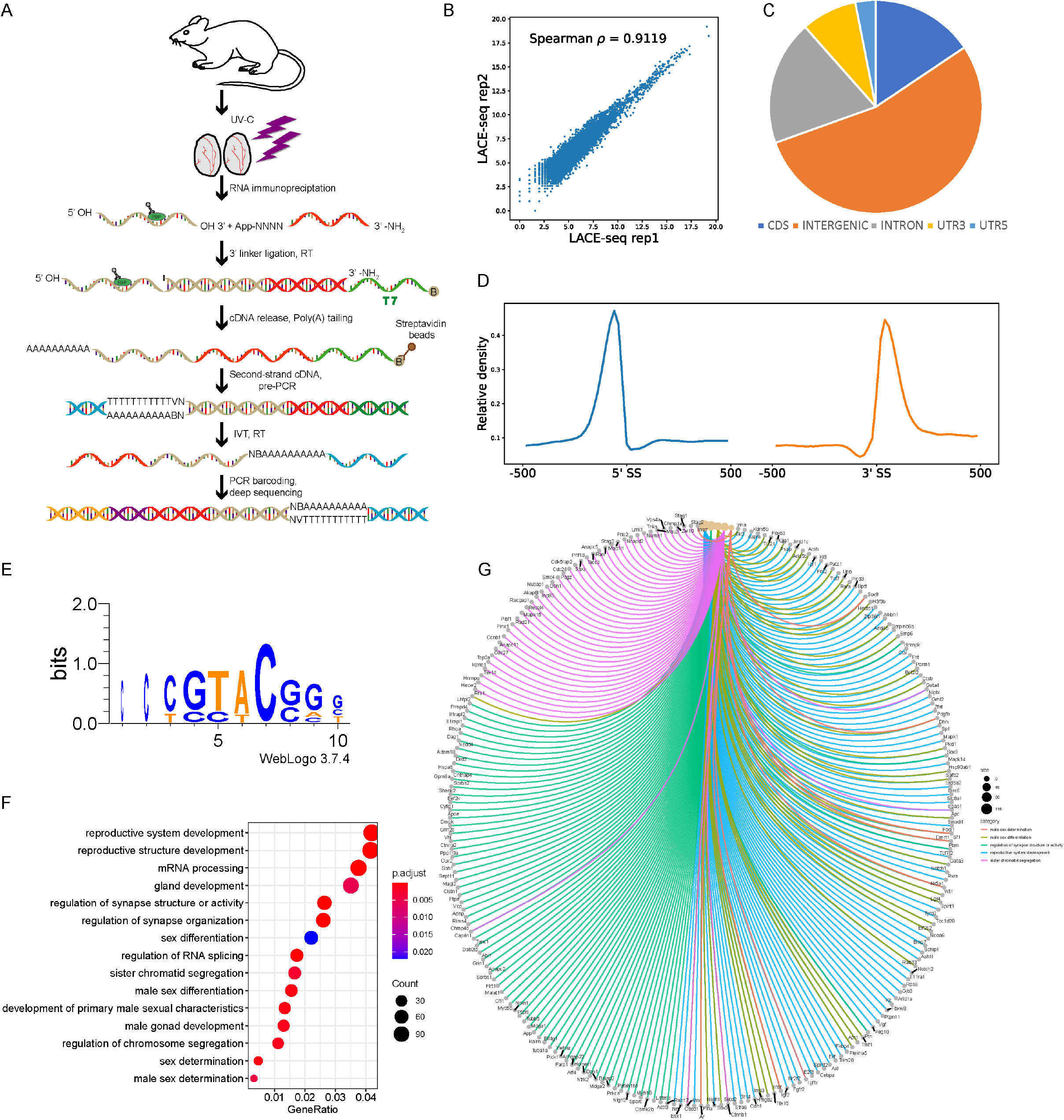
Global landscape of SRSF2-binding sites in mouse testes as revealed by using LACE-seq. (A) Flowchart of the LACE-seq method. RBP, represents RNA-binding protein. A circled B represents biotin modification. N, represents random nucleotide; V represents A, G or C. IVT, represents in vitro transcription. (B) Spearman correlation plot between SRSF2 LACE-seq replicates in total testes for assessing the reproducibility of the data. Spearman correlation for the reads counts of each sample was calculated from two replicates. (C) Genomic distribution of SRSF2 binding sites in testes. CDS, coding sequence. UTR3, 3′ untranslated region. UTR5, 5′ untranslated region. (D) Schematic analysis showing the distribution of SRSF2-binding sites in the vicinity of the 5′ exon-intron and the 3′ intron-exon boundaries (500 nt upstream and 500 nt downstream of 3′SS; 500 nt upstream and 500 nt downstream of 5′SS). (E) SRSF2-binding motifs identified by LACE-seq in mouse testes. (F) GO enrichment map of SRSF2-binding genes. (G) Network analysis of the enriched GO terms of SRSF2-specific targets.

### SRSF2 affects expression and AS of *Stra8, Stag3* and *Atr* in a direct manner

By combining RNA-seq data with LACE-seq identified peaks, we identified 262 downregulated, and 187 upregulated transcripts as direct targets of SRSF2 in testes (Supplementary file 3). To obtain more comprehensive information, similarly, we then performed GO annotation. GO analysis showed that both significantly upregulated genes and SRSF2-binding genes were involved in reproductive development, male sex differentiation, and germ cell development (Figure 6A and Figure 6B). And spermatogenesis, meiotic cell cycle, male gamete generation, chromosome segregation, DNA repair, and DNA recombination were significantly enriched among these both significantly downregulated genes and SRSF2-binding genes (Figure 6C and Figure 6D). We next validated these both significantly DEGs and SRSF2-binding genes which were involved in spermatogenesis by using quantitative polymerase chain reaction (qPCR) to check the mRNA abundance (Figure 6E and Figure 6F). These data reflected that deletion of SRSF2 directly affects the expression levels of critical genes involved in spermatogenesis.

**Figure 6.**
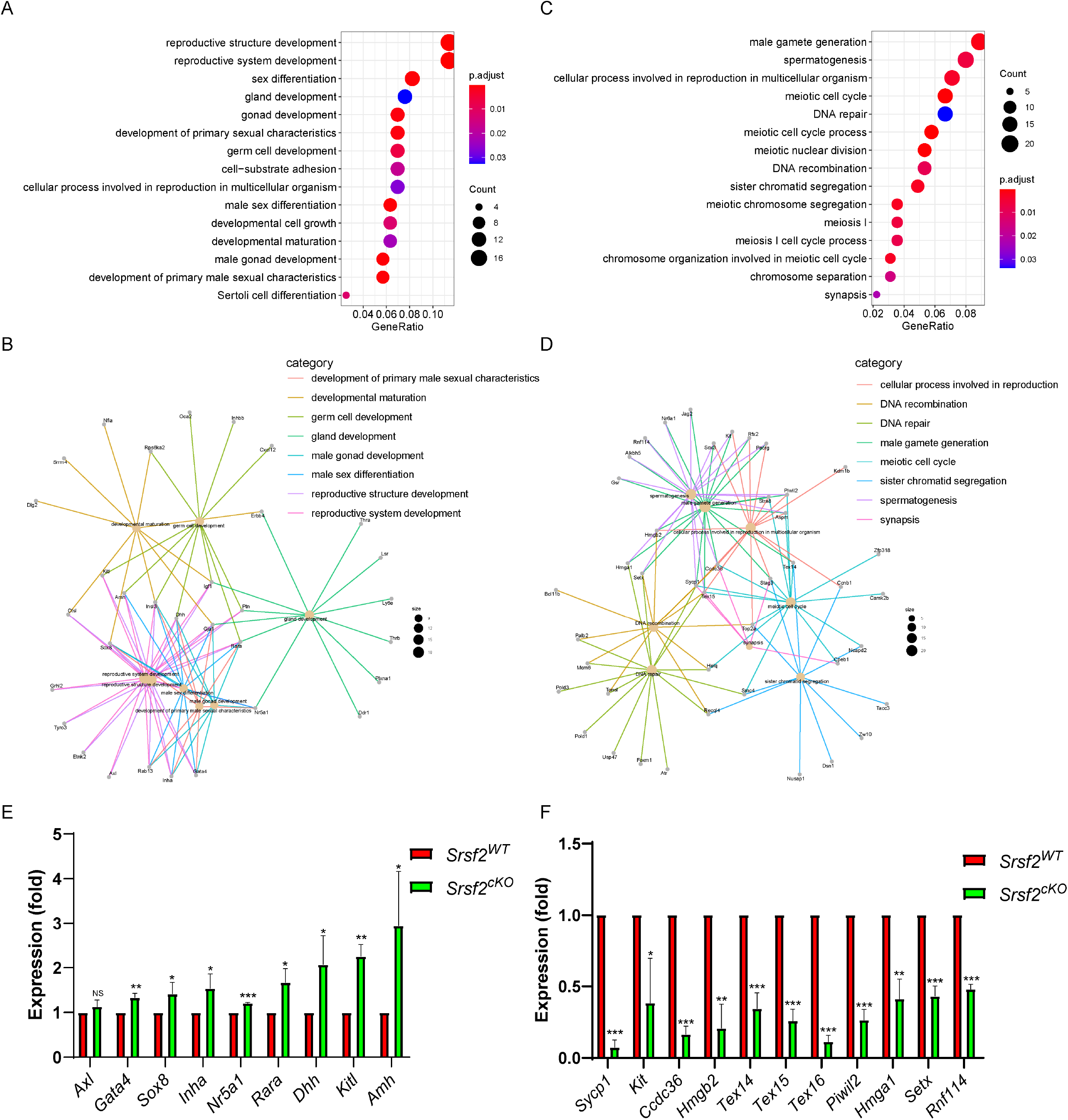
The expressions of key SRSF2-binding genes involved in the spermatogenesis change after *Srsf2* KO. (A) Correlation analysis between the RNA-seq and LACE-seq. GO analysis of the significantly upregulated genes and SRSF2-binding genes. (B) Network analysis of the enriched GO terms of the significantly upregulated genes and SRSF2-specific targets. (C) Correlation analysis between the RNA-seq and LACE-seq. GO analysis of the significantly downregulated genes and SRSF2-binding genes. (D) Network analysis of the enriched GO terms of the significantly downregulated genes and SRSF2-specific targets. (E) Quantitative RT-PCR validation of the expression of genes involved in (B). β -actin was used as the internal control. Data are presented as the mean ± SEM. *P*<0.05(*), 0.01(**) or 0.001(***). (F) Quantitative RT-PCR validation of the expression of genes involved in (D). β-actin was used as the internal control. Data are presented as the mean ± SEM. *P*<0.05(*), 0.01(**) or 0.001(***).

Furthermore, we investigated the relationship of SRSF2-binding genes, DEGs, and AS genes to confirm the direct targets that account for the abnormal spermatogenesis after SRSF2 cKO. Venn diagram showed that 14 SRSF2 directly binding genes were differentially down-regulated and spliced (Figure 7A). These genes included *Stra8, Stag3, Atr, Hmga1*, and *Setx*, and all of them were necessary for the male germ cell development (Figure 7B). We then researched SRSF2 regulatory mechanism on the expression of *Stra8, Stag3* and *Atr* by combining the RNA-seq with LACE-seq. The data showed that the abundance of *Stra8* mRNA was decreased and the ratio of exon 2 skipping was increased after SRSF2 cKO. Similarly, the abundance of *Stag3* mRNA was decreased and the ratio of exon 19 and 20 skipping was increased after SRSF2 cKO. The abundance of *Atr* mRNA was decreased and the ratio of exon 34 skipping was increased after SRSF2 cKO (Figure 7C). We also performed RT-PCR and semiquantitative reverse transcription PCR to confirm the above results (Figure 7D and Figure 7E). These experiments indicated that SRSF2 affects the expression levels and AS of *Stra8, Stag3* and *Atr* in a direct manner, which were critical for male germ cell differentiation and development.

**Figure 7.**
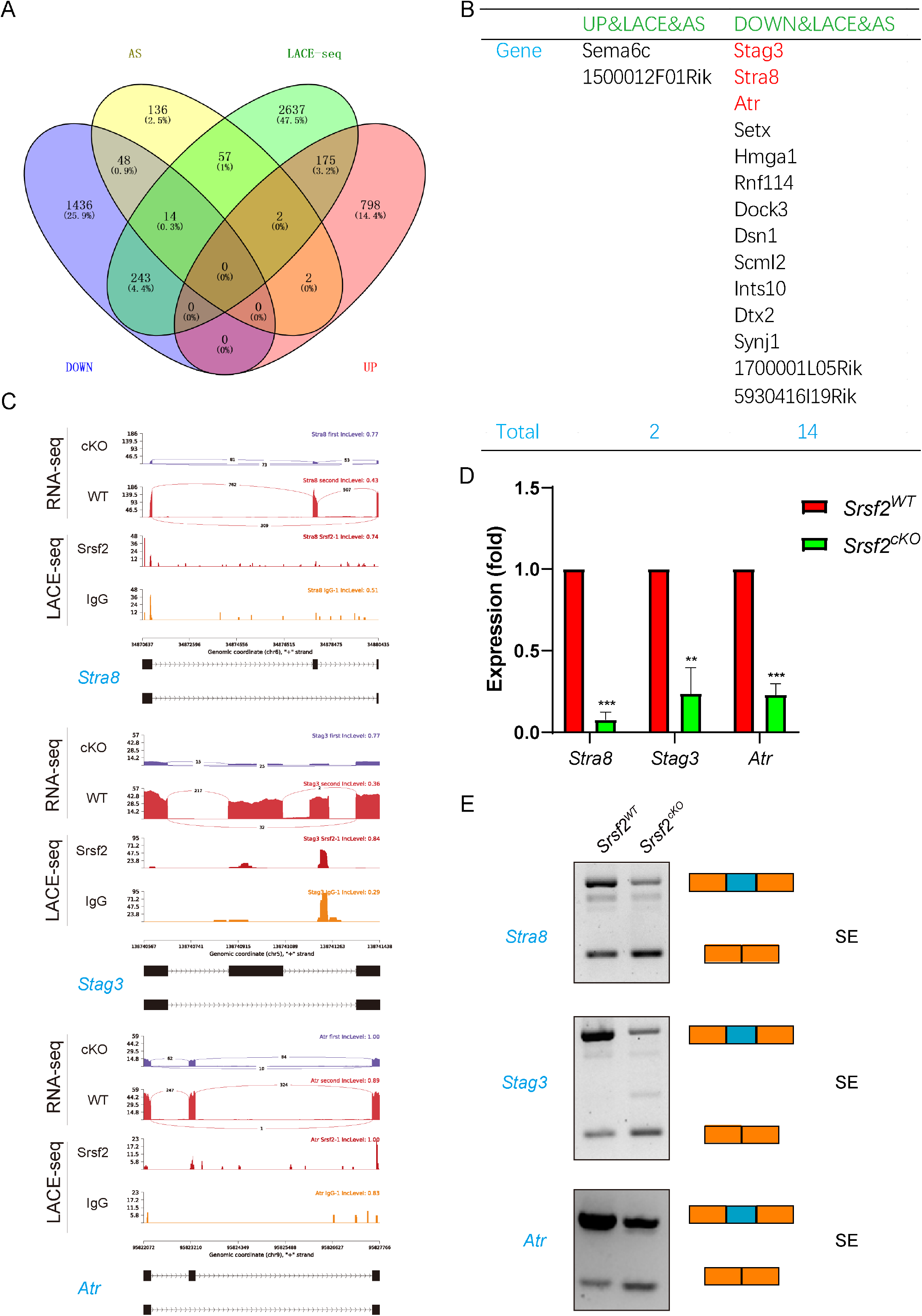
SRSF2 affects expression and alternative splicing of *Stra8, Stag3* and *Atr* in a direct manner. (A) Venn diagram shows the correlation among SRSF2-binding genes, DEGs, and AS genes. (B) The detailed genes of SRSF2-binding, differentially expressed, and AS. (C) A magnified view showing RNA-seq and LACE-seq signals of the selected candidate genes. IgG, immunoglobulin G. (D) Quantitative RT-PCR validation of the expression of *Stra8, Stag3*, and *Atr*. (E) Semiquantitative RT-PCR analysis of AS patterns of the changed spliced genes in *Srsf2*^*WT*^ and *Srsf2*^*cKO*^ testes at P10. PCR primers are listed in Supplementary file 1. The scheme and cumulative data on percentage of the indicated fragments are shown accordingly.

## Discussion

As members of the serine arginine-rich protein family, SRs which include 12 members in mammalian (SRSF1–12) are well-known for their regulatory function of splicing(32). The first SRs identified were SRSF1 (previously known as SF2/ASF) and SRSF2 (previously known as SC35) (33). SRs consist of one or two RNA-recognition motifs (RRM) in the N-terminus and arginine/serine amino acid sequences (RS domain) in the C-terminus(34). In general, RRM can recognize RNA and determine the binding of SRs to RNA, while the RS domain can regulate diverse protein-RNA and protein-protein interactions(33). Like other SR splicing factors, several studies in recent years have suggested that SRSF2 have important roles in regulating gene transcription, mRNA stability, genomic stability, and translation (22-25). Also, some findings suggested that SRSF2 may serve as a therapeutic target for various diseases (26-29).

Recently, it has been found that RNA-binding proteins (RBPs) have important functions during germline and early embryo development. As a RBP, SRSF2 is also expressed in testis, however, its functions in male germ cells is still completely unknown. In this study, by crossing *Srsf2*^*F/F*^ mice with *Stra8-Cre* mice to generate mutant mice, we found that SRSF2 is essential for spermatogenesis and fertility in males.

The RBPs could serve post-transcriptional functions to determine cellular RNA and protein levels. For the past few years, high throughput sequencing techniques have become an increasingly essential tool for biological research. RNA immunoprecipitation with sequencing (RIP-seq) and crosslinking immunoprecipitation coupled with high-throughput sequencing (CLIP-seq or HITS-CLIP) are two major methods to identify RBPs targets from millions of cells(35, 36). There are also some modified versions, such as iCLIP, irCLIP and eCLIP(37-39). Up to now, LACE-seq is the latest method developed by us, which can unbiasedly map the binding sites of these RBPs at single-nucleotide resolution in low-input cells (40). To gain a comprehensive perspective of the mechanisms of SRSF2 depletion in male germ cells, we isolated testes from wildtype mouse at P10 and systematically profiled binding landscape of SRSF2 proteins by using LACE-seq. The results showed that SRSF2 proteins could bind numerous genes in a direct manner. Then, our analysis showed that these SRSF2-binding genes were closely involved in the regulation of RNA splicing, reproductive development, male sex differentiation, regulation of synapse organization, and regulation of chromosome segregation. In addition, RNA-seq analysis further showed that transcriptome and splicing of transcripts change in SRSF2-null testes. By combining RNA-seq and LACE-seq data, we identified 262 downregulated, and 187 upregulated transcripts as direct targets of SRSF2 in testes. The two omics data reflected that deletion of SRSF2 directly affects the expression levels of critical genes involved in spermatogenesis, such as *Sycp1, Rnf114, Setx, Hmgb2, Gata4, Sox8, Amh, Kitl*, and *Axl*.

Retinoic acid (RA) is an important factor of spermatogenesis, with functions on spermatogonial differentiation and subsequently initiation of meiosis (41, 42). The two certain targets for RA are Stra8 and Kit. Several surveys indicated that *Stra8* has two different roles during spermatogenesis. On one hand, under the influence of RA, *Stra8* functions as a transcriptional repressor of the pluripotency program during differentiation of spermatogonia. When differentiating spermatogonia are near the end of their mitotic phase, *Stra8* switches to the second role and acts as a transcription activator of genes involved in meiosis initiation(43-45). In addition to RA signaling, *Dazl* is also regarded as a regulator of meiotic initiation(46). Of particular note, in *Srsf2*^*cKO*^ mice, the differentiation of spermatogonia and meiosis initiation were disrupted. Except for *Stra8, Stag3* and *Atr* are crucial regulators of meiotic processes during spermatogenesis(47-51). The two omics data also indicated that SRSF2 affects the expression levels and AS of *Stra8, Stag3* and *Atr* in a direct manner, which are critical for the male germ cell development process. Also, we found that the reduced expression and abnormal AS of *Dazl* were indirectly cuased by SRSF2 deletion (Figure 7-figure supplement 1A, B, C).

In summary, our study has demonstrated for the first time that SRSF2 has important functions in male fertility and spermatogenesis, especially in the differentiation of spermatogonia and meiosis initiation. Mechanistic analyses reveal that SRSF2 is essential for posttranscriptional regulation by specifically adjusting the gene expression and AS in direct or indirect manners during spermatogenesis. These abnormally expressed genes, such as *Stra8, Stag3, Atr* and *Dazl*, caused by SRSF2 deletion finally result in the failure of spermatogenesis and male infertility.

## Materials and Methods

### Mice

Mice lacking *Srsf2* in male germ cells (referred to as *Srsf2*^*cKO*^) were generated by crossing *Srsf2*^*Floxed/Floxed*^ (*Srsf2*^*F/F*^) mice with *Stra8-Cre* mice. All transgenic mouse lines have C57BL/6J genomic background. Genotyping PCR for *Srsf2* was performed using the following primers: forward: GTTATTTGGCCAAGAATCACA, and reverse: TAGCCAGTTGCTTGTTCCAA. The PCR conditions were as follows: 94 ℃ for 5 min; 35 rounds of 94 ℃ for 30 sec, 60 ℃ for 30 sec, and 72 ℃ for 30 sec; and 72 ℃ for 5 min. Genotyping PCR for *Stra8-Cre* was performed using the following primers: forward: ACTCCAAGCACTGGGCAGAA, wildtype reverse: GCCACCATAGCAGCATCAAA and reverse: CGTTTACGTCGCCGTCCAG. The PCR conditions were as follows: 94 ℃ for 5 min; 35 rounds of 94 ℃ for 30 sec, 60 ℃ for 30 sec, and 72 ℃ for 30 sec; and 72 ℃ for 5 min. Four genotypes in the progeny, including *Srsf2*^*F/+*^, *Srsf2*^*F/-*^, *Srsf2*^*F/+*^; *Stra8-Cre* and *Srsf2*^*F/-*^; *Stra8-Cre* were identified. The *Srsf2*^*F/+*^ male mice were used as control group. The mice were maintained under specific-pathogen-free (SPF) conditions and housed under controlled environmental conditions with free access to water and food. All animal operations were approved by the Animal Care and Use Committee of the Institute of Zoology, Chinese Academy of Sciences (CAS).

### Antibodies

β-actin antibody (mouse, sc-47778; Santa Cruz); SYCP3 (mouse, sc-74569; Santa Cruz); γH2AX (rabbit, 9718; Cell Signaling Technology, Inc.); MVH (mouse, ab27591; Abcam); SOX9 antibody (rabbit, AB5535, Sigma-Aldrich); PLZF antibody (goat, AF2944, R&D Systems); SFRS2 polyclonal antibody (rabbit, 20371-1-AP, Proteintech); SC35 antibody (mouse, S4045, Sigma-Aldrich); green-fluorescent Alexa Fluor® 488 conjugate of lectin PNA (L21409, Thermo). Horseradish peroxidase–conjugated secondary antibodies were purchased from Zhongshan Golden Bridge Biotechnology Co, LTD (Beijing). Alexa Fluor 488–conjugated antibody, 594–conjugated antibody and Alexa Fluor 647–conjugated antibody were purchased from Life Technologies.

### Breeding assay

Males of different genotypes (8 weeks) were used for the breeding assay. Each male mouse was caged with two wild-type ICR (Institute of Cancer Research) females (7 weeks), and their vaginal plugs were checked every morning. The number of pups in each cage was counted within a week of birth. Each male underwent six cycles of the above breeding assay.

### Immunoblotting

To prepare protein extracts, testes were homogenized in RIPA lysis buffer supplemented with protease and phosphatase inhibitor cocktail (Roche Diagnostics). After transient ultrasound treatment, the testis lysates were incubated on ice for 30 min and then centrifuged at 4 ℃, 12000 rpm for 20 min. The supernatant was transferred to a new tube and quantified using a BCA reagent kit (Beyotime, P0012-1). Then equal volume loading buffer was added. After being boiled at 95 ℃ for 10 min, the protein lysates were used for immunoblotting analysis. Immunoblotting was performed as described previously(52). Briefly, the separated proteins in SDS-PAGE were electrically transferred to a polyvinylidene fluoride membrane. After incubation with primary and secondary antibodies, the membranes were scanned with Bio-Rad ChemiDoc XRS+.

### Tissue collection and histological analysis

For histological analysis, at least three adult mice for each genotype were analyzed. Testes and caudal epididymides were dissected immediately following euthanasia. The tissues were then fixed in Bouin’s fixative (saturated picric acid: 37% formaldehyde: glacial acetic acid= 15: 5: 1) overnight at room temperature, dehydrated in an ethanol series, and embedded in paraffin wax. Then, 5μm sections were cut with a microtome. After 48 ℃ overnight drying, the sections were deparaffinized in xylene, hydrated by a graded alcohol series and stained with Hematoxylin and Eosin for histological analysis. Images were collected with a Nikon inverted microscope with a charge coupled device (CCD) (Nikon, Eclipse Ti-S, Tokyo, Japan).

### Immunofluorescence

Testes used for immunostaining were fixed in 4% paraformaldehyde (pH 7.4) overnight at 4 ℃, dehydrated, and embedded in paraffin. Paraffin-embedded testes were cut into sections of 5μm thickness. Then, the sections were deparaffinized, immersed in sodium citrate buffer (pH 6.0) and heated for 15 min in a microwave for antigen retrieval. After blocking with 5% donkey serum albumin, sections were incubated with primary antibodies at 4 ℃ overnight. Then the sections were incubated with an appropriate FITC-conjugated secondary antibody. The nuclei were stained with DAPI. Images were captured using a laser scanning confocal microscope LSM880 (Carl Zeiss, Germany).

### RNA extraction and gene expression analysis

Total RNA was extracted from whole testes using TRNzol Universal Reagent (cat. # DP424, Tiangen, China) according to the manufacturer’s instructions. Then reverse transcription (RT) was performed using the 5X All-In-One RT MasterMix (cat. # G490, Abm, Canada). RT-PCR was performed using the UltraSYBR Mixture (cat. # CW0957, Cowin Bio, China) on a LightCycler 480 instrument (Roche). The results were analyzed based on the 2^-^ΔΔCt method to calculate the fold changes. *β-actin* was used as an internal control. At least three independent experiments were analyzed. All primer sequences are listed in the Supplementary file 1. Semiquantitative PCR experiment was carried out with primers (listed in Supplementary file 1) amplifying endogenous transcripts. Then the PCR products were detected on 2% agarose gels. *Gapdh* was used as an internal control.

### RNA sequencing and data analysis

Total testes samples were used from P10 *Srsf2*^*WT*^ and *Srsf2*^*cKO*^ male mice according to three individual collections. One Total RNA was extracted from whole testes using TRNzol Universal Reagent (cat. # DP424, Tiangen, China) according to the manufacturer’s instructions. The quality of RNA samples was examined by NanoDrop 2000&8000 and Agilent 2100 Bioanalyzer, Agilent RNA 6000 Nano Kit. The high-quality RNAs were used to prepare the libraries, followed by high-throughput sequencing on an Illumina NovaSeq 6000. The RNA sequencing experiment was supported by Annoroad BioLabs.

After trimming adaptor sequence and rRNA, the retained reads from *Srsf2* control and cKO samples were aligned to mouse genome (mm9) using HISAT2 with default parameters. Only non-RCR duplicate and uniquely mapped reads were used for subsequent analysis. Significantly changed genes were screened using DESeq2 with |log2FC| > 1 and FDR<0.05. Alternative splicing events were identified by rMATS with default parameters. Only events with FDR<0.001 and splicing difference > 0.05 were regarded as significant.

### LACE-sequencing and data analysis

Total testes samples were used from P10 WT male mice for LACE-seq. LACE-seq method was performed as described recently by us (40). Briefly, the samples were firstly irradiated twice with UV-C light on ice at 400 mJ. Then RNA immunoprecipitation of the samples was performed. The immunoprecipitated RNAs were then fragmented by MNase and dephosphorylated. Then a series of steps were performed to include, reverse transcription, first-strand cDNA capture by streptavidin beads, poly(A) tailing, pre-PCR, IVT, RNA purification, RT, PCR barcoding and deep sequencing.

The adapter sequences and poly(A) tails at the 3′ end of raw reads were removed using Cutadapt (v.1.15) with two parameters: -f fastq -q 30,0 -a ATCTCGTATGCCGTCTTCTGCTT -m 18 --max-n 0.25 --trim-n., and -f fastq -a A -m 18 -n 2. Clean reads were first aligned to mouse pre-rRNA using Bowtie, and the remaining unmapped reads were then aligned to the human (hg19) or mouse (mm9) reference genome. For LACE-seq data mapping, two mismatches were allowed (Bowtie parameters: -v 2 -m 10 --best -strata; -v 2 -k 10 --best -strata). Peaks were identified by Piranha with parameters: -s -b 20 -p 0.01. Peaks without IgG signal were selected for further usage. For motif analysis, LACE-seq peaks/clusters were first extended 30 nt to 5′ upstream, and overrepresented hexamers in the extended sequences were identified as previously described(53). The consensus motifs were generated from the top-10 enriched hexamers using WebLogo.

## Statistical analysis

All of the experiments were performed at least three times independently. Paired two-tailed Student’s t-test was used for statistical analysis. Data analyses were carried out via GraphPad Prism 8.00 (GraphPad Software, Inc.) and presented as mean ± SEM and *P*<0.05(*), 0.01(**) or 0.001(***) was considered statistically significant.

## Data availability

The data sets from this study have been submitted to the NCBI Gene Expression Omnibus (GEO; https://www.ncbi.nlm.nih.gov/geo/) under accession number: GSE 206537.

## Acknowledgements

We thank Dr Xiang-Dong Fu for providing the *Srsf2*^*F/F*^ mice. We appreciate and acknowledge Shiwen Li and Xili Zhu for their technical assistance. We thank all members of the Sun lab for their help and discussion. This study was supported by National Key R&D Program of China (2018YFA0107701, 2019YFA0109900), Guangdong Basic and Applied Basic Research Foundation (2021A1515111118) and the Shenzhen High-level Hospital Construction Fund.

## Conflict of Interest

The authors declare no conflict of interest.

## Figure Legends

**Figure 3-figure supplement 1.**
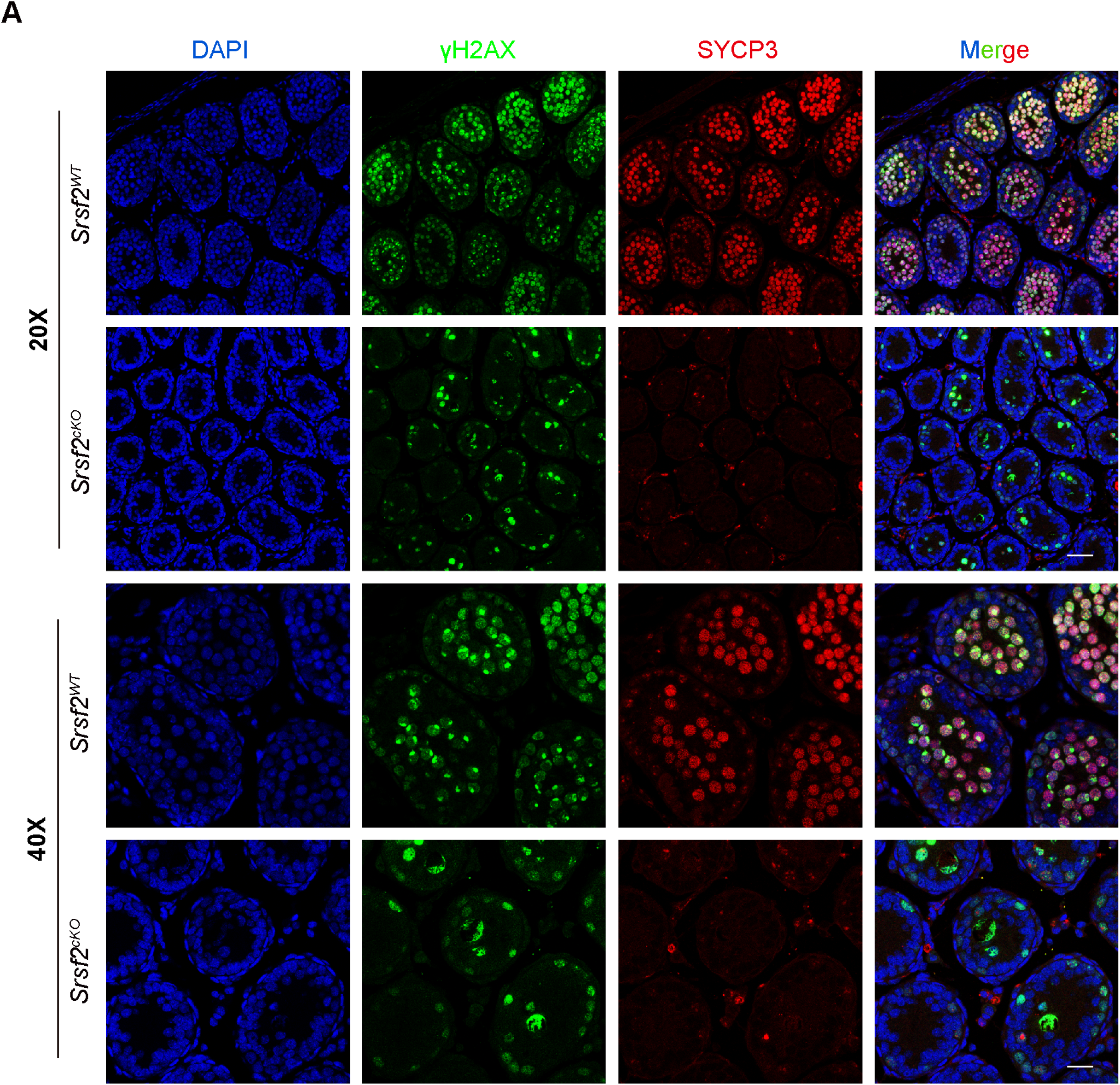
Spermatogenesis fails to progress into meiosis in *Srsf2* deficient germ cells at P12. γH2AX (green) and SYCP3 (red) immunofluorescence analysis of the *Srsf2*^*WT*^and *Srsf2*^*cKO*^ male mice at P12. Scale bar: (top) 50 μm; (bottom) 20 μm.

**Figure 4-figure supplement 1.**
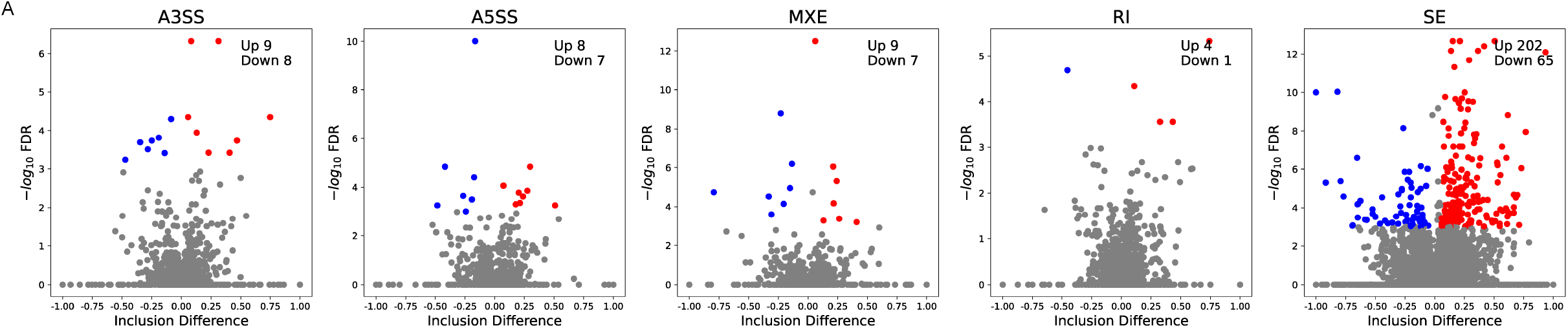
SRSF2 regulates mRNA alternative splicing in testes. Five AS events significantly affected by deletion of SRSF2 in the testes at P10. The different types of alternatively spliced events were shown.

**Figure 7-figure supplement 1.**
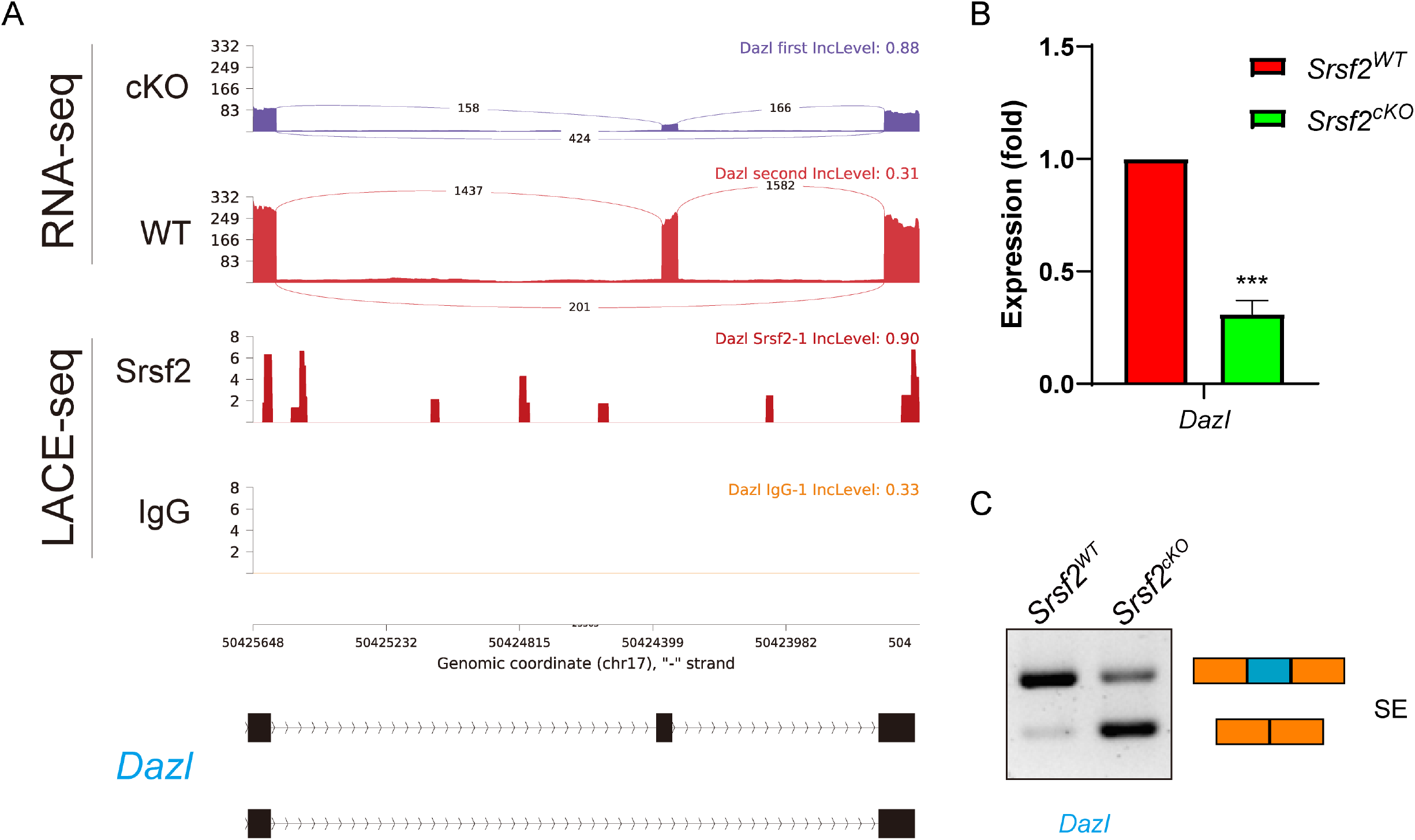
SRSF2 indirectly regulates splicing and expression of *Dazl*. (A) A magnified view showing RNA-seq signals of the *Dazl* gene. (B) Quantitative RT-PCR validation of the expression of *Dazl*. (C) Semiquantitative RT-PCR analysis of AS patterns of the 732 changed spliced genes in *Srsf2*^*WT*^ and *Srsf2*^*cKO*^ testes at P10. PCR primers are listed in Supplementary file 1. The scheme and cumulative data on percentage of the indicated fragment are shown accordingly.

**Figure 1-source data 1 Actin and SRSF2 protein levels**

**Figure 1-source data 2 The fertility of *Srsf2***_***cKO***_ **male mice**.

**Figure 2-source data 1 The testis weight of adult male mice.**

**Figure 6 source data 1 Quantitative RT-PCR validation of the expression**

**of genes**.

**Figure 7-source data 1 Quantitative RT-PCR validation of the expression**

**of *Stra8, Stag3*, and *Atr***.

**Figure 7-source data 2 Semiquantitative RT-PCR analysis of AS patterns of the changed spliced genes**.

**Figure 7-figure supplement 1-source data1 Quantitative RT-PCR**

**validation of the expression of *Dazl***.

**Figure 7-figure supplement 1-source data2 Semiquantitative RT-PCR analysis of AS patterns of the changed spliced genes**.

## Notes

### Competing Interest Statement

The authors have declared no competing interest.

